# Pirin does not bind to p65 or regulate NFκB-dependent gene expression but does modulate cellular quercetin levels

**DOI:** 10.1101/2024.12.03.626411

**Authors:** Melissa Meschkewitz, Erika M Lisabeth, A. Denaly Cab-Gomez, Jeffrey Leipprandt, Richard R Neubig

## Abstract

Pirin is a non-heme iron binding protein with a variety of proposed functions including serving as a co-activator of p65 NFκB and quercetinase activity. We report here, failure to confirm pirin’s primary proposed mechanism, binding of Fe(III)-pirin and p65. Analytical size exclusion chromatography (SEC) and fluorescence polarization (FP) studies did not detect an interaction. We also found no effects of pirin on TNFα-activated p65-regulated gene transcription using mouse embryonic fibroblasts (MEFs) from a pirin knockout mouse and a pirin knockdown NIH3T3 fibroblast cell line. TNFα - activated p65 response gene mRNA was neither increased nor decreased in cells with loss of pirin compared to wildtype cells. Furthermore, pirin immunofluorescence in NIH3T3 fibroblasts showed primarily a cytoplasmic localization, not nuclear as in most previous studies. This was confirmed by cell fractionation analysis. Pirin did show colocalization with the endoplasmic reticulum (ER) marker protein disulfide-isomerase (PDI) as well as cyotoplasmic labeling. We confirmed pirin’s quercetinase activity in biochemical assays and demonstrated competitive inhibition by the pirin inhibitor CCG-257081. Cellular quercetin levels in cells exposed to quercetin *in vitro* were increased by knockdown of pirin or by treatment with pirin inhibitors. Since pirin is localized to ER and flavanols are protective of ER stress, we investigated whether pirin knockdown altered ER stress signaling but did not find any effect of pirin knockdown on ER stress response genes. Our results challenge the dominant model of pirin’s function (NFκB regulation) but confirm its quercetinase activity with implications for the mechanisms of pirin binding small molecules.

**Significance Statement:** Pirin has multiple proposed functions and plays an important role in cancer (melanoma, colon, and breast) and inflammatory diseases. Small molecule pirin-binding compounds have been identified but pirin’s functional mechanism remains poorly understood. We raise doubts about the primary description of pirin as a nuclear regulator of p65 NFκB function but validate pirin’s role as a quercetinase. We show that pirin-binding compounds can raise cellular quercetin levels. Further studies will be required to fully understand pirin’s biological mechanisms.

## Introduction

Pirin was first discovered as a potential co-activator of nuclear factor I/CCAAT box transcription factor (NFI/CTF1)(Wendler *et al*., 1997). Structural analysis of pirin revealed that it is a member of the cupin superfamily of proteins in plants, which have a conserved beta barrel fold. Pirin is comprised of two beta barrel domains and coordinates iron in the cavity of its N-terminal domain as a non-heme iron-binding protein (Pang *et al*., 2004). Pirin is highly conserved in mammals, plants, fungi, and prokaryotes, and shares structural similarity with a related protein YhhW from *Escherichia coli*. Enzymatic analysis of human pirin and YhhW shows that both possess quercetinase activity. They also degrade other flavonols with high affinity (Adams and Jia, 2005; Guo *et al*., 2019).

Pirin’s functions in mammalian cells, however, are relatively poorly understood. It seems to play a key role in cancers (Arai *et al*., n.d.; Wang *et al*., 2014; Perez-Dominguez *et al*., 2021) and potentially in inflammatory processes (Brzóska *et al*., 2014; Hu *et al*., 2021; Zhang *et al*., 2022; Shen *et al*., 2024). As its most frequently cited function, pirin has been proposed to be a redox-sensitive regulator of the p65 subunit of NF-κB (Liu *et al*., 2013; Adeniran and Hamelberg, 2017; Talà *et al*., 2018). Pirin was reported to interact with the Rel Homology Domain of p65 in surface plasmon resonance (SPR) studies and modeling suggested that Fe(III)-pirin can modulate binding of p65 to DNA (Liu *et al*., 2013; Adeniran and Hamelberg, 2017). Cell-based studies also found that the human papillomavirus protein E7 promotes NF-kB activation in a pirin-dependent manner (Carrillo-Beltrán *et al*., 2020). On the other hand, pirin has been modeled to interact with master regulators of inflammation and additional pathways, like BCL3, NFΚBIA, NFIX and SMAD9 in its Fe(II)-bound form (Dechend *et al*., 1999; Ahsan *et al*., 2023). Another cellular role for pirin has been found in autophagy-induced ferroptosis where it protects from DNA damage during ferroptosis; pirin downregulation increases DNA damage and leads to activation of autophagy-induced ferroptosis (Hu *et al*., 2021).

We reported that pirin binds to a series of small molecule compounds developed as inhibitors of Rho/MRTF/SRF signaling (Lisabeth *et al*., 2019). This CCG series of compounds, has anti-fibrotic, anti-metastatic and anti-resistance properties (Haak *et al*., 2014b, 2017; Hutchings *et al*., 2017; Yu-Wai-Man *et al*., 2017; Lionarons *et al*., 2019; Misek *et al*., 2020). Several other chemical inhibitors of pirin have also been identified (Miyazaki *et al*., 2010; Cheeseman *et al*., 2016) with effects on cancer cell function. Knockdown of pirin, as well as pharmacological inhibition with the pirin-binding CCG and CCT compounds, decreases SRF-dependent gene transcription in primary dermal fibroblasts (Lisabeth *et al*., 2019). This suggests another link between pirin and gene transcription.

Here, we investigate multiple roles for pirin with the primary aim of confirming the role of pirin in p65 NFκB-mediated gene transcription. The primary objective was to confirm pirin binding to p65/DNA and modulating p65 signaling and then investigate the effect of CCG compounds on pirin binding to p65. However, we were not able to detect Fe(III)-pirin-binding to p65/DNA and in fibroblasts find no evidence for pirin in the nucleus. To test for other effects of pirin on p65 signaling, we generated pirin knockout mice and a knockdown cell line in NIH3T3 fibroblasts. Again, we did not observe differences in p65 signaling in the absence of pirin. Therefore, we were unable to confirm a role for pirin in p65 signaling. We did confirm pirin’s quercetinase activity and show that pirin knockdown cells showed higher quercetin levels in the presence of exogenous quercetin. Also, the CCG and CCT pirin-binding compounds inhibit pirin’s enzymatic activity and increase cellular quercetin levels. Thus, a key proposed function of pirin on p65 NFκB signaling appears not to be correct and further studies will be required to fully understand pirin’s mechanisms in cancer and other mammalian cell functions.

## Materials and Methods

### Protein purification

Pirin: A His6-tagged pirin expression plasmid was developed and pirin protein purified from *E. coli* as previously described (Lisabeth *et al*., 2019) with the following modifications. To prepare apo-pirin and Fe(III) pirin, BL21 *E. coli* was grown in M9 minimal media (1x M9 salts, Sigma Aldrich, Burlington Massachusetts, #M6030-1Kg; 0.4% glucose, 2 mM MgSO_4_, 0.3 mM CaCl_2)_. After purification, his-tagged pirin was treated with 10 equivalents of EDTA in lysis buffer (50mM NaH_2_PO_4_ pH 8.0, 300mM NaCl, 5% glycerol) at 4°C overnight then subjected to size exclusion chromatography on a Superdex 200 column in 50 mM NaH_2_PO_4_, pH 8.0, 150mM NaCl then 5% glycerol was added (storage buffer) prior to storage at −80°C. To prepare Fe(III) pirin, apo-pirin was incubated overnight at 4°C with 1.2 equivalents of Fe(III) chloride hexahydrate and desalted on a PD SpinTrap^TM^ G-25 (GE Healthcare, Chicago, Illinois) into pirin storage buffer. Bovine serum albumin (BSA) for control experiments was treated in the same manner (treatment with Fe(III) and desalting).

P65 Rel Homology Region (P65RHR): The p65 RHR was amplified from 3xFLAG full length p65 (DNASU, with primers 5’-CTAGCTAGCATGCCCTATGTGGAGATCATTG-3’ (Fwd) and 5’-CGCGGATCCTTACCGGGGGTCGGTGGGT-3’ (Rev) by PCR and digested with restriction enzymes NheI and BamHI. The digested PCR product was then ligated into pET11a which had been digested with the same restriction enzymes prior to ligation. The P65 RHR (AA 19-325) expression plasmid was transformed into BL21 cells and cultured on LB Agar plates. One liter of LB with Ampicillin was inoculated and cells were grown at 37°C and shaking at 200 rpm until an OD_600_ of 0.1 was reached. Protein production was induced with 0.1mM IPTG for 16 hours at 22°C. Cells were pelleted by centrifugation at 6000 rpm (10,000xg) for 15 minutes and resuspended in 50 mL of lysis buffer (25 mM Tris-HCl pH 7.0, 50 mM NaCl, 0.5 mM EDTA, 10 mM β-Mercaptoethanol; β-Me) with protease and phosphatase inhibitors (Thermo Scientific, Waltham, Massachusetts, #A32961). Lysates were sonicated on ice (Cole Parmer ultrasonic homogenizer 4710) for 6 minutes (12 x 30 sec) and centrifuged at 12,000 rpm (24,515xg) for 40 minutes to clear. Cleared lysate was treated with 10% streptomycin sulfate (final concentration 1%) on ice for 20 minutes to precipitate nucleic acids and centrifuged again. After the second centrifugation, the lysate was filtered with 0.45 µm filter, and protein was purified on an SP Sepharose column in lysis buffer with a gradient elution of NaCl (50mM-500mM). Fractions containing protein were concentrated and further purified by size exclusion chromatography on a Superdex200 column and stored at −80°C (p65 storage buffer 25 mM Tris HCl pH 7.5, 50 mM NaCl, 1 mM DTT).

### Size Exclusion Chromatography Analysis of pirin and P65 binding

P65RHR and double stranded IgκB DNA (5’-GAGTTGAGGGGACTTTCCCAGGC-3’) were incubated at equimolar concentrations (50 µM) for 1 hour at room temperature in gel filtration buffer (25 mM Tris HCl pH 7.5, 50 mM NaCl) and unbound DNA was removed through gel filtration on a Superdex 200 size exclusion Column (GE Healthcare, Chicago, Illinois). The protein concentration of P65RHR and DNA complex was determined through BCA assay. P65RHR/DNA complex (20 µM) was then incubated with 40 µM of Fe (III)-pirin for 1 hour at room temperature before analysis for complex formation on Superdex 200 10/300 GL size exclusion column. Peak fractions (21-33) were collected separated by SDS-PAGE and protein was detected through Coomassie staining (p65 RHD) and Western Blotting (pirin). Membranes were blocked with Licor blocking buffer (Licor, #927-70001) and pirin was then detected on a LI-COR Odyssey CLx (Primary antibody: 1:1,000 dilution from Cell Signaling, Danvers, Massachusetts; Secondary Antibody: 1:10,000 dilution, Licor, Lincoln, NE, IRDeye^R^ 680RD Goat anti-Rat, #926-68076)

### Fluorescence Polarization Analysis of p65/DNA binding

FP was performed in black 384-well plates (Greiner, Kremsmuenster, Austria, Ref#781076) and polarization was measured on a BioTek Synergy Neo (485 nm excitation, 528 nm emission). For p65 binding curves, 5 nM annealed FAM-labeled IgκB DNA (5‘-56-FAM-GAGTTGAGGGGACTTTCCCAGGC-3’) was incubated with p65 concentrations varying from 3 nM to 4 µM for 1 hour at room temperature before measuring polarization. To test effects of pirin on p65 binding of DNA, p65 and DNA were incubated with 500 nM Fe(III)-pirin. Polarization was calculated by subtracting perpendicular emission from parallel emission, dividing by the sum of the emissions and multiplying by 100.

### Generation of regulated pirin knockdown (shPirin) and control (shVector) cell lines (Figure S1)

NIH 3T3 fibroblasts were purchased from Elabscience (Houston, Texas, #EP-CL-0171) and maintained in 10% New Born Calf Serum (NBCS; Gibco, Brooklyn, New York, #16010159), DMEM (Gibco, Brooklyn, New York, #11995-065) with Antibiotics/Antimitotics (A/A; Gibco, Brooklyn, New York, #15240-062) shRNA oligo design was done through the RNAiConsortium collection (MISSION shRNA, Sigma, Burlingtion, Massachusetts), and three shRNA sequences were chosen.

Forward and reverse oligos were purchased for each shRNA sequence and annealed before ligating into pTetOn pLKO-puro (a gift from Dmitri Wiederschain, Addgene plasmid #21915 (Wiederschain *et al*., 2009)) that was previously digested with AgeI and EcoRI. Positive clones were verified by sequencing.

#1:CCGGGATGTATATATTGGGCCTGATCTCGAGATCAGGCCCAATATATACATC TTTTTTG

#2:CCGGCTGGACTTCAAGTTGGACCAACTCGAGTTGGTCCAACTTGAAGTCCAGTTTTTTG

#3:CCGGGTTGTCCAACATGGTCCATTTCTCGAGAAATGGACCATGTTGGACAA CTTTTTTG

Virus particles were produced by transient transfection of HEK293T cells with 5000 ng of pTetOn pLKO vector containing shPirin #1, #2, #3 (or empty vector control), 500 ng of pMD2.G, and 5000 ng of psPAX2 with Lipofectamine 2000 (Thermo Scientific, Waltham, Massachusetts, #11668019). Four hours after transfection, medium was changed to DMEM containing 10% FBS (Gibco, Brooklyn, New York #10437-028). Twenty-four hours after transfection, medium containing lentivirus was collected and filtered through a 0.45 mm filter and frozen at −80.

NIH 3T3 cells (800,000 cells in a 10 cm plate) were infected with 6 mL of lentivirus particles and 10 µl of 10 mg/mL polybrene. Twenty-four hours after infection, the medium was changed. Then 48 hours after infection, the cells were split to a 15-cm plate and the virally transduced cells were selected with 2 µg/mL puromycin. After all cells from a non-infected control culture died, selection was considered complete (72 hours total). shRNA expression was induced by adding 100 ng/mL doxycycline for 48 hours. Knockdown was determined by analysis of pirin mRNA by qPCR and protein by western blot. shPirin #2 was the most effective at reducing pirin mRNA and protein (Figure S1) and was used for subsequent studies.

### Generation of pirin knockout mice

Pirin knockout mice were generated by the MSU Transgenic and Genome Editing Facility. All animal procedures complied with the US National Institutes of Health guidelines on animal care and were approved by the MSU Institutional Committee on Animal Care and Use.

C57B6 mice were used as founders for knockout mouse development. A frameshift was created by introducing a single nucleotide A in exon 2 of the pirin gene in E18 embryos. The guide RNA sequence 5’-TCAGCCGGGAGCAGTCAGAG GGG-3’ was used for RNP electroporation targeting in C57B6 zygotes. Successful founders were determined by DNA sequence analysis. The founders were backcrossed at least 4 times with C57B6/J wildtype mice. Since the pirin gene is on the X chromosome, male mice carrying the mutant allele (Pir^X-/Y^) are full knockouts while females may be heterozygous (Pir^-/+^) or homozygous (Pir^-/-^).

### Genotyping of mice

Genotyping was performed by restriction fragment analysis of PCR products containing the target site. Primers 5’-TCCATCTTCTACAGCTCAACAT-3’ and 5’-TGGCCTTCTCTCTCCAAATCC-3’ were used to amplify genomic DNA using 32 cycles of denaturation (95°C for 30 sec.), annealing (55°C for 30 sec.), and elongation (72°C for 30 sec.). The 235 bp PCR product was digested with restriction enzyme Hpy188III (New England Biolabs, Ipswich, Massachusetts, cat# R0622S), The WT sequence remains uncut, whereas the KO product is digested into 137bp and 99bp fragments.

### Generation of Primary Mouse Embryonic Fibroblasts (MEFs)

Pregnant female mice, approximately E13.5 were euthanized and mouse embryonic fibroblasts (MEFs) were generated as described in (Isolation of Mouse Embryo Fibroblasts (Durkin *et al*., n.d.). Briefly, embryos were trypsinized with 2.5% Trypsin-EDTA (Gibco, Brooklyn, New York, #25200-056) and dispersed cells plated in DMEM supplemented with 10% FBS and 1% A/A. Embryos were genotyped and both male and female MEFs were used for experiments. Cells were passaged approximately once per week and used within 10 passages.

### Gene expression analysis by qRT-PCR

#### NFκB activation

shPirin and shVector cell lines were grown in DMEM supplemented with 10% NBCS and 1% A/A. Each line was split into two flasks (with/without 100 ng/ml doxycycline) for 72 hours. Cells were then plated in 6-well plates at 400,000 cells/well. Cells induced with doxycycline 72 h prior were kept in medium with doxycycline. The next day, cells were washed with PBS and the medium changed to DMEM containing 0.5% NBCS with no doxycycline added. After 24 hours, cells were stimulated with 10 ng/ml TNFα for 1 hour. Following incubation, RNA was isolated using a Zymo RNA isolation kit for RNA (ZymoGenetics, Seattle, Washington, #R1055) and cDNA was made using approximately 1000 ng of RNA with a High-Capacity cDNA Reverse Transcription Kit (Thermo Fisher Scientific, Waltham, Massachusetts, #4368814). qPCR was done using a SYBR green Master mix kit (Thermo Scientific, Waltham, Massachusetts, #4039155) using primers against GAPDH, pirin, IL-6, and IκBα (sequences below). The ΔΔCt method was used for calculating fold-expression changes. Values in each experiment were normalized to those for unstimulated shVector cells.

#### ER stress activation

shPirin and shVector cell lines were grown and induced as described above. Cells were plated in 6-well plates at 400,000 cells/well, 72 h after induction with doxycycline. The next day, cells were stimulated for 5 hours using Tunicamycin or Thapsigargin at a concentration of 5 µg/ml and 1 µM, respectively. RNA was then isolated and cDNA made as described above. qRTPCR and analysis of results was done as described above, using primers against GAPDH, pirin, CHOP, ATF4, sXBP1, usXBP1, totalXBP1, BiP, GRP94, and EDEM. Results were normalized to those for unstimulated shVector cells.

### qPCR primers

GAPDH Fwd: 5’-TGACCTCAACTACATGGTCTACA-3’

GAPDH Rev: 5‘-CTTCCCATTCTCGGCCTTG-3’

pirin Fwd: 5‘-GTCGAAGGTTTACACTCGCAC-3’

pirin Rev: 5‘-AATAGGCTGGGAATGCTTTGC-3’

IL-6 Fwd: 5‘-CTGCAAGAGACTTCCATCCAG-3’

IL-6 Rev: 5‘-AGTGGTATAGACAGGTCTGTTGG-3’

IκBα Fwd: 5‘-TGAAGGACGAGGAGTACGAGC-3’

IκBα Rev: 5‘-TGCAGGAACGAGTCTCCGT-3’

CHOP Fwd: 5’-CCACCACACCTGAAAGCAGAA-3’

CHOP Rev: 5’-AGGTGAAAGGCAGGGACTCA-3’

ATF4 Fwd: 5’-GGGTTCTGTCTTCCACTCCA-3’

ATF4 Rev: 5’-AAGCAGCAGAGTCAGGCTTTC-3’

sXBP1 Fwd: 5’-CTGAGTCCGAATCAGGTGCAG-3’

sXBP1 Rev: 5’-GTCCATGGGAAGATGTTCTGG-3’

usXBP1 Fwd: 5’-CAGCACTCAGACTATGTGCA-3’

usXBP1 Rev: 5’-GTCCATGGGAAGATGTTCTGG-3’

total XBP1 Fwd: 5’-TGGCCGGGTCTGCTGAGTCCG-3’

total XBP1 Rev: 5’-GTCCATGGGAAGATGTTCTGG-3’

BiP Fwd: 5’-TTCAGCCAATTATCAGCAAACTCT-3’

BiP Rev: 5’-TTTTCTGATGTATCCTCTTCACCAGT-3’

GRP94 Fwd: 5’-AAGAATGAAGGAAAAACAGGACAAAA-3’

GRP94 Rev: 5’-CAAATGGAGAAGATTCCGCC-3’

EDEM Fwd: 5’-CTACCTGCGAAGAGGCCG-3’

EDEM Rev: 5’-GTTCATGAGCTGCCCACTGA-3’

### Quercetinase Assays

The quercetinase assay was performed as described (Guo *et al*., 2019). Briefly, pirin was diluted to 5 µM in 50 mM NaPO_4_, pH 8.0, 150 mM NaCl, 5% glycerol. The assay was performed with 2.5 µM pirin in 200 µl final volume with 50 mM NaPO_4_, pH 7.4, 150 mM NaCl, and compounds or DMSO control were added to pirin at room temperature, 30’ before the addition of quercetin. Absorbance was recorded every 30 seconds for 5’ at 380 nm in a 96-well UV-transparent microplate (Corning, Corning, New York, #3679) and read in a Biotek Synergy Neo at room temperature. Rates of change were calculated by fitting the data to a linear regression model (GraphPad Prism). CCG-257081 was obtained from Dr. Scott Larsen at the University of Michigan (Ann Arbor, MI).

### Cellular quercetin levels with 2-aminoethyl diphenylborinate (APB) Fluorescence

An APB fluorescence assay was used to detect cellular quercetin levels (Rozanski *et al*., 2019). After 72h of incubation with doxycycline, shPirin and shVector cells were plated in black wall/clear bottom 384-well plates (Greiner, Kremsmuenster, Austria, #781091) at 4,000 cells/well in DMEM containing 10% NBCS and 1% A/A. After 24 h, quercetin at the indicated concentrations was added to cells for 40 min. Media were removed and cells washed with 1x PBS. The cells were then incubated with either APB in PBS (62.5 µg/ml) or kept in PBS for 10 min. After incubation, APB/PBS or PBS was removed and cells were lysed in 20 µl lysis buffer (250 µg/ml APB, 0.35% Triton-X in PBS) and then read for fluorescence (excitation at 485 nm, emission at 550 nm) on a Biotek Synergy Neo plate reader. For treatment with compounds, 3T3 cells were plated in a similar manner. Cells were treated with CCG257081 or CCT251236 overnight before addition of quercetin. As a control, cells without exogenous quercetin were read for APB fluorescence which showed very low endogenous levels of flavonols.

### Western Blotting

Cells were lysed in RIPA buffer supplemented with protease and phosphatase inhibitors (Pierce by Thermo Scientific, Waltham, Massachusetts, #A32961) and sonicated for 10 seconds using a microtip sonicator (Fisher Scientific Sonic dismembrator model 500). Lysates were then spun at max speed in a tabletop centrifuge at 4^0^C for 10 minutes and the protein concentration of the supernatant was determined by BCA assay (Pierce BCA Protein Assay, by Thermo Scientific, Waltham, Massachusetts, #23228). Approximately 50 µg of total protein was loaded onto a 12.5% Polyacrylamide gel and transferred for 2 hours at 140V to a PVDF membrane. Membranes were blocked with LiCor blocking buffer (#927-70001) and then probed using the appropriate antibodies (see below).

### Nuclear and Cytoplasmic Fractionation

Approximately 1×10^6^ cells were plated in a 6-cm plate in 10% FBS+DMEM supplemented with A/A. Cells were treated with TGF-β (Research and Diagnostics Systems, Inc., Minneapolis, Minnesota), and then 200 µl of fractionation buffer (20 mM Hepes, pH 7.4, 10 mM KCl, 2 mM MgCl2, 1 mM EDTA, 1 mM EGTA with protease and phosphatase inhibitors Thermo Scientific, Waltham, Massachusetts, #A32961 and 1 mM DTT) and was added to the cells. Cells were scraped and lysates were kept on ice for 15 minutes, passed through a 25-gauge needle 10 times, and kept on ice for an additional 20 minutes. Nuclei were pelleted by centrifugation at 3,000 rpm (735 x g) and washed twice with fractionation buffer and lysed in 100 µl RIPA buffer. Cytosolic fractions were then concentrated to the same volume as that of the nuclear fractions using spin columns (Millipore, Burlington, Massachusetts, #UFC501096) and equal volumes of the two samples were loaded onto SDS-PAGE gels for analysis by western blotting.

### Immunocytochemistry

NIH 3T3 fibroblasts were grown in DMEM supplemented with 10% NBCS and 1% A/A. Before the experiment, glass coverslips were sterilized in 70% EtOH and coated with 10 µg/ml fibronectin (R&D Systems, Minneapolis, Minnesota, #1030-FN) in PBS for 1 hour at room temperature. Cells were plated on coverslips in 12-well plates at 20,000 cells/well and incubated overnight at 37^0^C. The next day, the medium was changed to DMEM containing 0.5% NBCS. Twenty-four hours after the medium change, cells were stimulated with 10 ng/ml TNFα (R&D, #210-TA) for 1 hour. Between each subsequent step, cells were washed 3x with 1xPBS. Cells were fixed in 4% paraformaldehyde in PBS (Biotum, Fremont, California, #22023) for 10 minutes at room temperature and permeabilized with 0.25% Triton-X-100 in PBS for 10 minutes. Cells were then treated with Image-iT FX signal enhancer (Invitrogen by Thermo Scientific, Waltham, Massachusetts, #R37107) for 30 minutes at 37°C. Cells were then blocked with 10% goat serum (Vector Labs, Newark, California, #S-1000-200) in PBS with a 1:1000 dilution of Triton-X-100 for 1 hour at room temperature. Primary antibody (rat anti-pirin(1E8), Cell Signaling, Danvers, Massachusetts, #9777S) was added to cells overnight at 4°C at a dilution of 1:100 in 2% BSA with 1:1000 Triton-X-100. The next day, the cells were incubated with a 1:1000 dilution of secondary antibody (AlexaFluor^TM^ 488 goat anti-rat, Invitrogen by Thermo Scientific, Waltham Massachusetts, #A11006) in 2% BSA with 1:1000 dilution of Triton-X-100 at room temperature for 1 hour. Coverslips were then mounted with ProLong Gold antifade reagent with DAPI (Invitrogen by Thermo Scientific, Waltham, Massachusetts, #R36935) on glass microscopy slides or stained with rhodamine phalloidin (Cytoskeleton Inc, Denver, Colorado, #PHDR1) following the manufacturer’s protocol before mounting. After drying for 24 hours, slides were imaged using an Eclipse Ti2-E Inverted Motorized Research Microscope System – Nikon.

#### Pirin and ER colocalization

The following adaptations were made to the above protocol. The day after cells were plated on coverslips in 12-well plates, cells were washed with warm 1x PBS and fixation was performed using 4% formaldehyde solution/methanol-free (Thermo Scientific, Waltham, Massachusetts, #28906) for 15 min at 37 C. Cells were washed 3x with 1x PBS and permeabilized with 0.2% Triton-X-100 in PBS for 10 minutes at room temperature. Samples were treated with Image-iT™ FX Signal Enhancer (Invitrogen by Thermo Scientific, Waltham, Massachusetts, #R37107) for 30 min at 37C. Blocking was carried out using 2% BSA with a 1:1000 Triton-X-100 in PBS for 1 hr at room temperature. Cells were treated overnight with protein disulfide isomerase (PDI) monoclonal antibody RL90 (#MA3-019) and pirin (1E8) Rat Monoclonal Antibody (Cell Signaling, Danvers, Massachusetts, #9777) at dilutions of 1:75 for PDI and 1:100 pirin in 2% BSA. Secondary antibody treatment was performed using Alexa Fuor-488 goat-anti rat IgG (Invitrogen by Thermo Scientific, Waltham, Massachusetts, #A11006) and Alexa Fluor-568 donkey anti-mouse IgG (H+L) (Invitrogen by Thermo Scientific, Waltham, Massachusetts, #A10037) for 1 hr at room temperature, protected from light. Coverslips were then mounted and imaged as described above.

### Statistical Analysis

Statistical analysis was performed using GraphPad Prism 10. Specific methods are detailed in figure legends including one-way ANOVA or 2-way ANOVA with the indicated post-tests to correct for multiple comparisons. The quercetinase enzymatic activity was analyzed using the Extra Sum of Squares F-test in GraphPad Prism to test the null hypothesis that CCG-257081 did not inhibit the quercetinase activity.

## Results

### Recombinant pirin does not bind to p65 *in vitro*

We attempted to confirm the published (Liu *et al*., 2013) surface plasmon resonance (SPR) results for pirin binding to P65RHR bound to biotinylated IgκB DNA on an SPR chip, but we did not see a significant concentration-dependent difference in SPR-signal between p65 alone (50 nM) and with pirin (12.5 - 200 nM) (Figure S2. Therefore, we tried alternative approaches to confirm pirin binding to the NFκB p65/DNA complex. As in SPR, we used the purified Rel Homology Region (RHR) of p65 complexed with IgκB promoter DNA in combination with Fe(III)-pirin (Figure 1). Size-exclusion chromatography analysis of a mixture of p65RHR/DNA with Fe(III)-pirin at 40 µM concentrations did not show any evidence of new peak representing a stable complex or a shift in the migration of pirin to higher apparent molecular weight (Figure 1A). The p65RHR/DNA peak alone eluted at fraction 24 (apparent MW ∼100 kDa) and the Fe(III)-pirin alone eluted at fraction 31 (apparent MW ∼20 kDa). The mixture of p65RHR/DNA and Fe (III)-pirin resulted in a chromatogram that was identical to the summation of the chromatograms for the individual proteins alone. The identity of the peaks and location of p65 and pirin were confirmed by western blot and Coomassie staining (Figure 1B).

**Figure 1.**
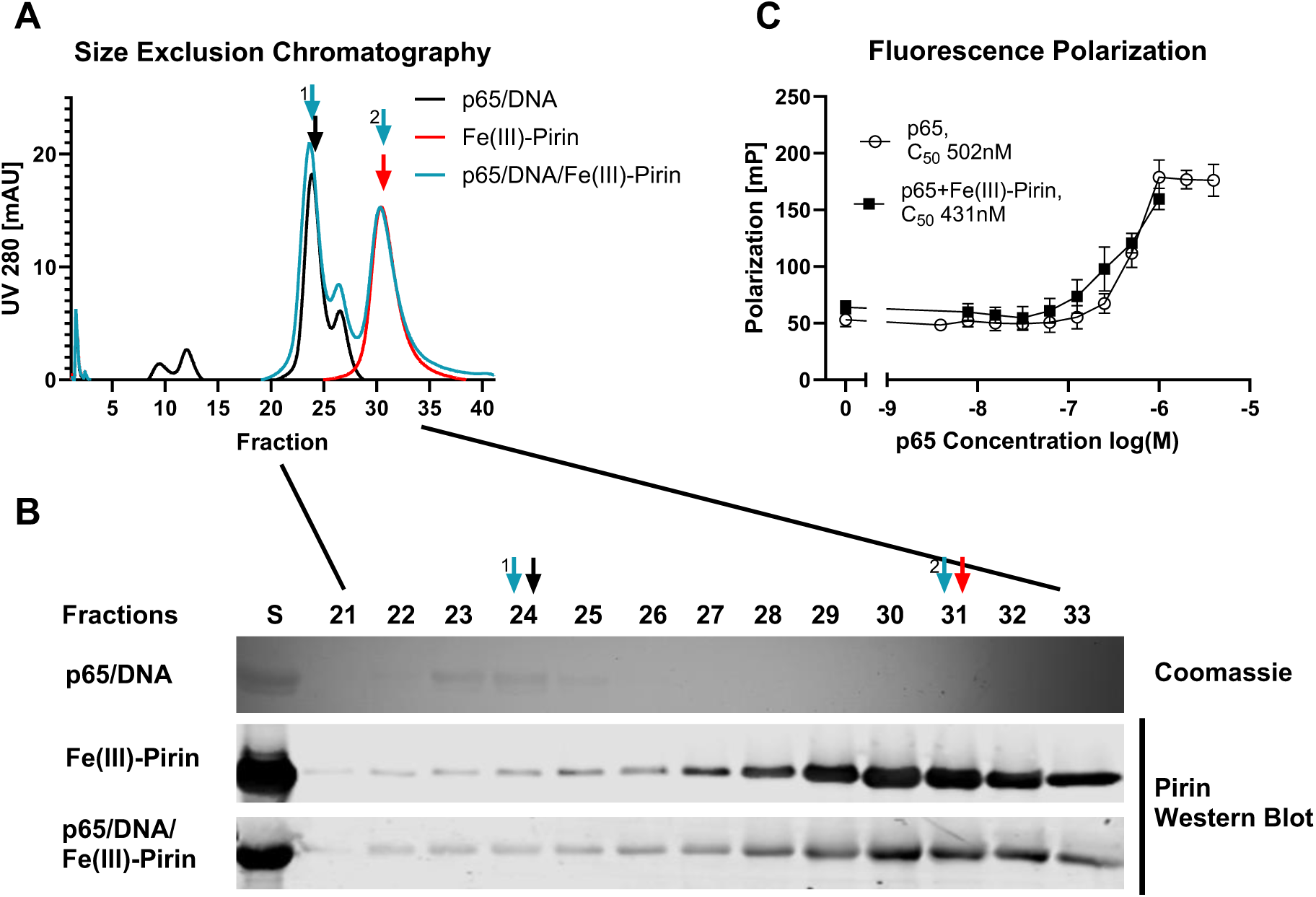
Binding of Fe (III)pirin to p65. (A) Gel filtration of 20 µM p65 with DNA (black), 40 µM Fe(III) pirin alone (red), and p65 with DNA together with Fe (III) pirin (light blue). The curves are representative of n=3. Fractions of 0.5 ml were collected from gel filtration and samples run on a 12.5% acrylamide gel (B) top lane Coomassie staining for p65 and bottom lanes Western Blot for pirin. Western blotting for p65 could not be done due to nonspecific binding of p65 Rel Homology Domain antibodies. S represents the sample before it was subjected to gel filtration. (C) Fluorescence polarization showing p65 binding curve with or without Fe(III)pirin. Error bars show standard deviation of n=3 biological replicates.

As a third method, we asked whether pirin could stabilize p65RHR binding to DNA by a fluorescence polarization analysis with FAM-labeled kappaB DNA. Increasing concentrations of p65RHR (3 – 4,000 nM) added to a fixed amount of DNA (5 nM) resulted in a 3.5-fold increase in polarization with a concentration of 500 nm P65RHR producing a half-maximal increase in polarization (Figure 1C). Addition of Fe(III)-pirin (500 nM) did not significantly increase the magnitude of polarization or change the concentration of p65 at which the half-maximal increase was achieved (430 nM). (Figure 1C). Overall, our biochemical experiments do not support an interaction between pirin and p65RHR.

### Pirin is localized in the cytoplasm and endoplasmic reticulum rather than in the nucleus in fibroblasts

If pirin did bind to the p65/DNA complex, one might expect to find pirin localized in the nucleus. Indeed, pirin was first reported to be a nuclear protein (Wendler *et al*., 1997). However, since that initial report, several publications have found that pirin is localized in the cytoplasm rather than in the nucleus (Licciulli *et al*., 2010; Ma *et al*., 2024). Given the importance of the pirin-binding compounds such as CCG-257081 in fibrosis and myofibroblast activation, we explored the cellular localization of pirin in fibroblasts with and without stimulation by ligands involved in inflammation and fibrosis (TNFα and TGF-β, Figure 2). Pirin is excluded from the nucleus with a cytoplasmic and perinuclear distribution in NIH 3T3 cells (Figure 2A). The distribution by immunocytochemistry does not change when cells were stimulated with TNFα. Cells were stained for F-actin to visualize cell structure and margin. The perinuclear pattern of pirin suggested to us possible endoplasmic reticulum (ER) localization. Using protein disulfide isomerase (PDI) as a marker for the ER, we found that there is substantial overlap of pirin staining with that of the ER marker (Figure 2B).

**Figure 2.**
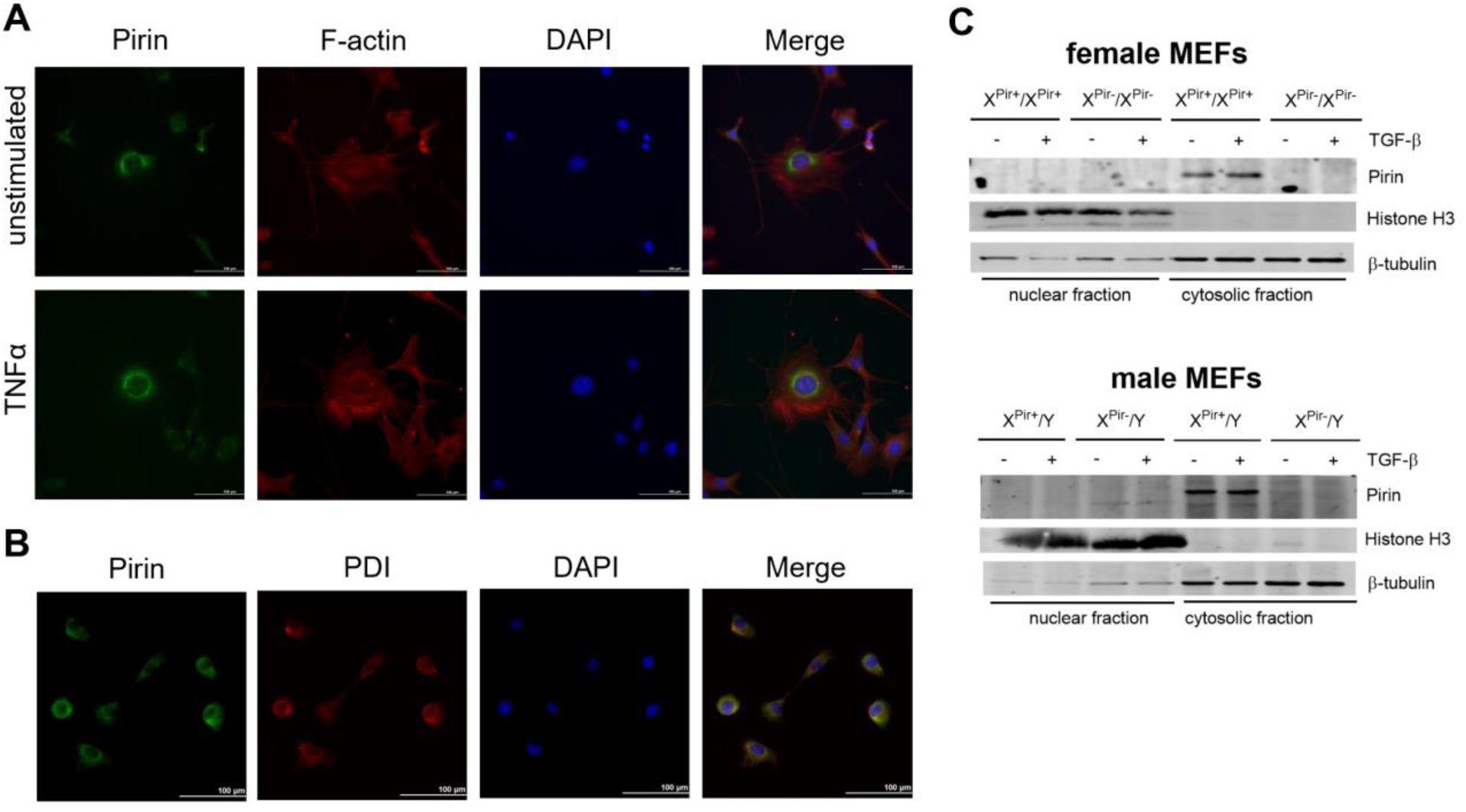
Pirin localization. (A) Immunofluorescence with antibody against pirin (FITC) and F-actin staining in 3T3 cells unstimulated or stimulated with TNFα. Nuclei are stained with DAPI. Images are representative of n=3. (B) Immunofluorescence with antibodies against pirin (FITC) and PDI (TRITC) as ER marker. (C) Nuclear fractionation and western blots from MEFs from male and female wild type and knockout mice probed for pirin. Histone H3 was used as marker for the nuclear fraction, β-tubulin was used as marker for cytosolic fraction.

We also used biochemical fractionation and Western blotting to assess pirin’s subcellular localization in MEFs generated from wildtype and pirin knockout mice (Figure 2C). Histone H3 was used as a nuclear marker and β-tubulin as a cytosolic marker. MEFs generated from female and male knockout mice did not show the immunostained band at 32 kDa seen in WT fibroblasts. This supports the specificity of the antibody and confirms the completeness of the knockout. Wildtype mice showed pirin staining only in the cytosolic fractions with no signal detected in the nuclear fractions. With our fractionation methodology (735 x g for 5 minutes), ER would be collected with the cytosolic fraction. TGF-β stimulation of the MEFs, which indirectly activates Rho and MRTF signaling (Ji *et al*., 2021), also did not result in pirin translocation to the nuclear fraction.

### Modulation of pirin expression does not alter NFκB-dependent gene expression

While we were not able to detect pirin-binding to NFκB p65 in biochemical assays, it is possible that pirin could interact in the cellular context and serve as a regulator of NFκB-mediated gene transcription in cells as previously proposed (Liu *et al*., 2013; Ma *et al*., 2024). To assess the effects of pirin on NFκB-mediated gene expression, we generated a regulated pirin knockdown NIH 3T3 fibroblast cell line with a pirin-specific small hairpin RNA (shRNA – shPirin cells) and control cells with vector alone and no pirin shRNA (shVector). Treatment of the shPirin but not shVector cells with doxycycline suppressed pirin mRNA levels by 66% (Figure 3C, Figure S1; 81.8% for Figure S1) and protein by ∼70% (Figure S1). Using these regulated pirin knockdown cells, we determined the effects of pirin on expression of NFκB response genes. After pretreatment with doxycycline for 72 hours, cells were treated with 10 ng/ml TNFα to activate the nuclear translocation of NFκB and induce expression of the NFκB-dependent genes IL-6 and IκBα (Vanden Berghe *et al*., 1998; Hayden and Ghosh, 2014; Liu *et al*., 2017). We saw similar TNFα induction of IL-6 and IκBα gene expression in shPirin and shVector cell lines regardless of treatment with doxycycline or not (Figure 3A & B, Figure S3). When treated with lower concentrations of TNFα we saw a slight increase in the expression of IL-6 when comparing shVector cells treated with doxycycline to shPirin cells treated with doxycycline (Figure S3E). However, at 0.1 ng/ml no change in gene expression for IL-6 was seen and no change was observed for expression of IκBα mRNA with either concentration (Figure S3E&F).

**Figure 3.**
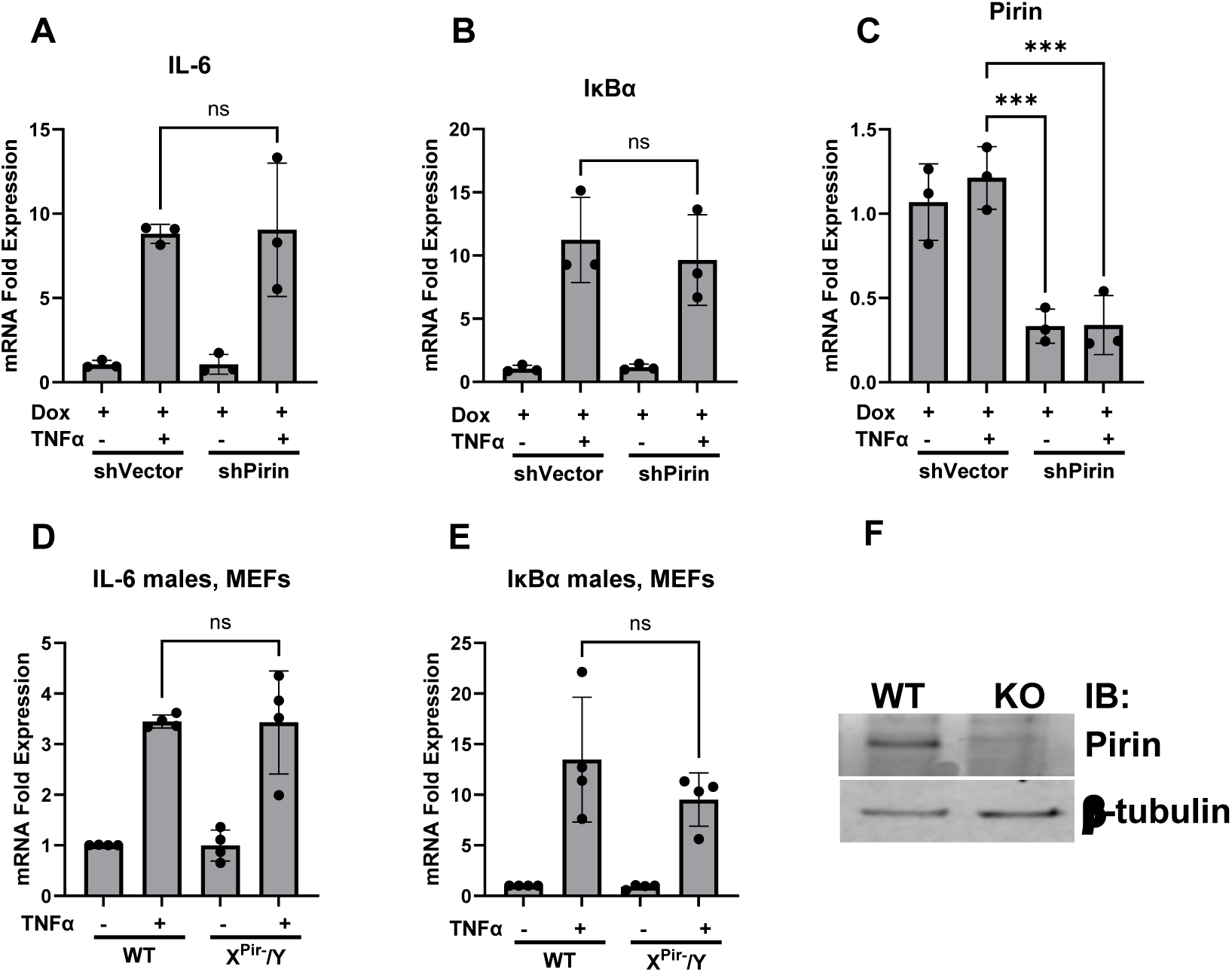
Expression of NFκB response genes. Fold-expression change of mRNA of IL-6 (A), IκBα (B) and Pirin (C) of stable transfected inducible pirin knockdown 3T3 fibroblasts (shP) (n=3). Cells were stimulated with TNFα to induce IL-6 and IκBα expression. Stably transfected 3T3 cells with vector DNA (shVector) were used as control. Cells were stimulated with doxycycline (+Dox) to induce pirin knockdown in shPirin cells. Fold-expression of mRNA was determined with RT-qPCR and the ΔΔCt method and expression was normalized to shVector cells without doxycycline or TNFα. (D&E) Mouse Embryonic Fibroblasts (MEFs) generated from male wildtype and pirin knockout mice were stimulated with TNFα and expression of IL-6 mRNA (D) and IκBα mRNA (E) was measured and analyzed as described in above (n=4). (F) Western Blot of fibroblasts from mice to confirm knockout is shown on the right. Statistical analysis was done by One-way ANOVA in GraphPad Prism 10 with significance values of * = p<0.05, ** = p<0.01, and *** = p<0.001. Error bars show standard deviation.

Similarly, in Mouse Embryonic Fibroblasts (MEFs) generated from male hemizygous pirin knockout or wildtype mice (see breeding outcomes Figure S6), we saw the same TNFα induced increases in IL-6 and IκBα gene expression in wildtype and knockout cells (Figure 3D & E). Global knockout of pirin in our knockout mice was confirmed with Western blots (Figure S4).

### Pirin is a quercetinase and its enzymatic activity is inhibited by pirin-binding compounds

Pirin has been reported to have 2,3 dioxygenase activity against eight different flavonols, including quercetin and kaempferol (Adams and Jia, 2005; Guo *et al*., 2019). However, it is not known if this activity has any effect in a cellular context or whether it can be modulated by pirin-binding compounds such as CCT 251236 or CCG-257081 (Cheeseman *et al*., 2016; Lisabeth *et al*., 2019). Using an *in vitro* enzymatic assay based on absorbance at 380 nm, pirin has a K_m_ of 18.8 µM for quercetin as the substrate (Figure 4A). The pirin-binding compound CCG-257081 (Lisabeth *et al*., 2019) reduced the degradation of quercetin in a concentration-dependent manner. Fitting the entire data set with a competitive inhibition model provided a good fit with a K_i_ value for CCG-257081 of 11.5 µM (Figure 4A).

**Figure 4.**
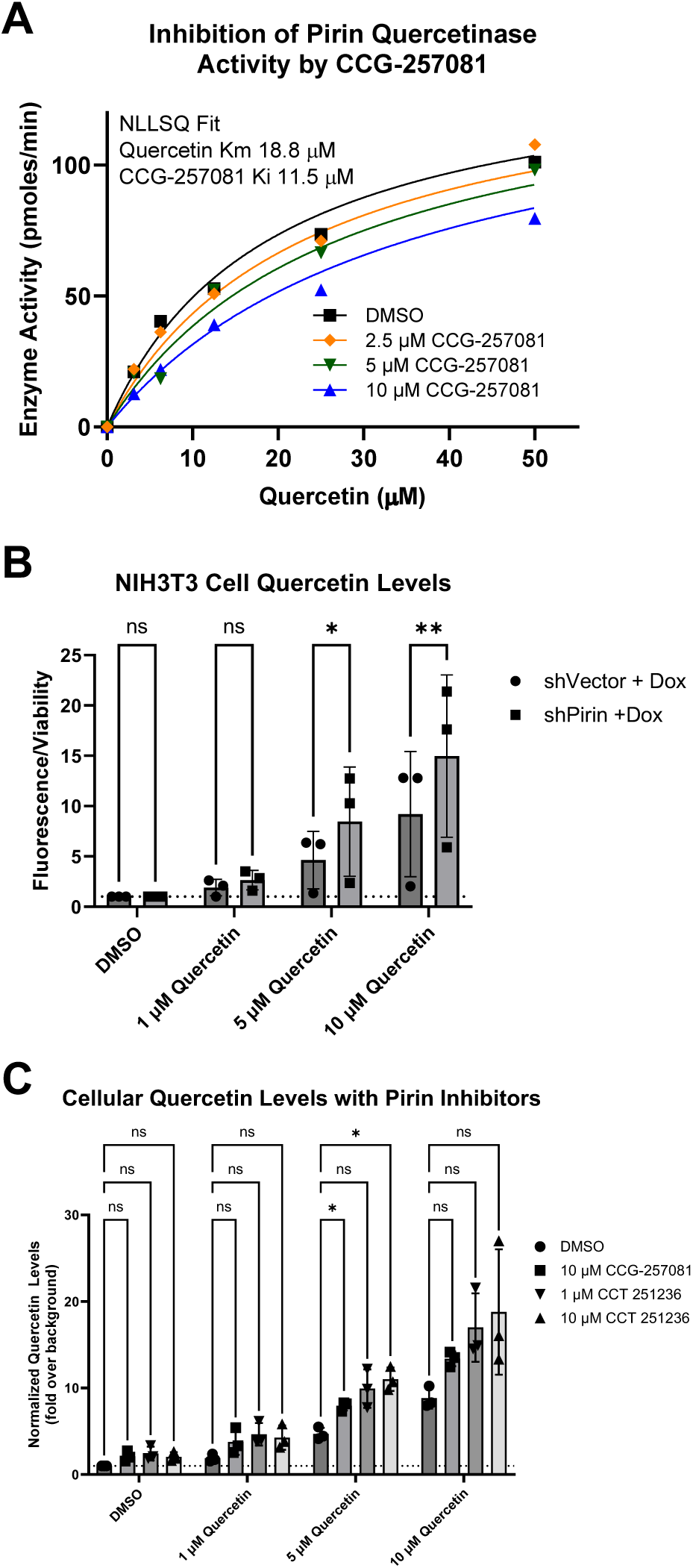
Pirin metabolism of quercetin. (A) *In vitro* degradation of quercetin by pirin with inhibition of pirin by CCG-257081. Statistical analysis was done by the Extra Sum of Squares F-test based on the null hypothesis that CCG-257081 did not inhibit the enzyme activity, ***, p<0.001 (B) Levels of exogenous quercetin in shVectorector and shPirin cells. (C) Inhibition of quercetin degradation in 3T3 cells by pirin inhibitor compounds. Cellular quercetin levels in B and C were measured using 2-aminoethyl diphenylborinate (APB) and normalized to viability measured with Celltiter Glo. Statistical analysis used 2-way ANOVA with a Sidak (B) or Dunnet (C) post-test to determine the significance of individual comparisons. Significance values are represented by * = p<0.05, ** = p<0.01, and *** = p<0.001.

Next, we used our doxycycline-inducible shPirin cells to investigate whether modulation of pirin levels in cells could affect quercetin levels in the presence of exogenous quercetin. Additionally, we tested whether pirin inhibitors (e.g. CCG-257081) could modulate quercetin levels in cells (Figure 4B & C). shPirin cells in the presence of doxycycline had higher levels of cellular quercetin than did the shVector cells. Thus, pirin does appear to degrade quercetin in a cellular context (Figure 4B). Next, we tested quercetin levels in NIH 3T3 fibroblasts after addition of compounds known to bind pirin, including our compound CCG-257081(Hutchings *et al*., 2017; Lisabeth *et al*., 2019) and CCT251236 (Cheeseman *et al*., 2016). Addition of these pirin-binding compounds increased exogenous quercetin levels about 1.5-2-fold with the addition of 1 μM quercetin (Figure 4C). This is consistent with these compounds inhibiting the quercetinase activity of pirin in cells.

### Pirin knockdown does not affect ER stress-induced gene expression

Since pirin shows significant co-localization with ER, we investigated a potential role for pirin in ER stress induction. Using tunicamycin and thapsigargin, we induced ER stress in doxycycline-treated shPirin and shVector cells, and measured gene expression levels of a panel of ER stress genes (CHOP, ATF4, sXBP1, usXBP1, total XBP1, BiP, GRP94, EDEM). There were no significant differences in ER stress gene expression between NIH-3T3 cells with and without pirin knockdown (Figure 5, Figure S5).

**Figure 5.**
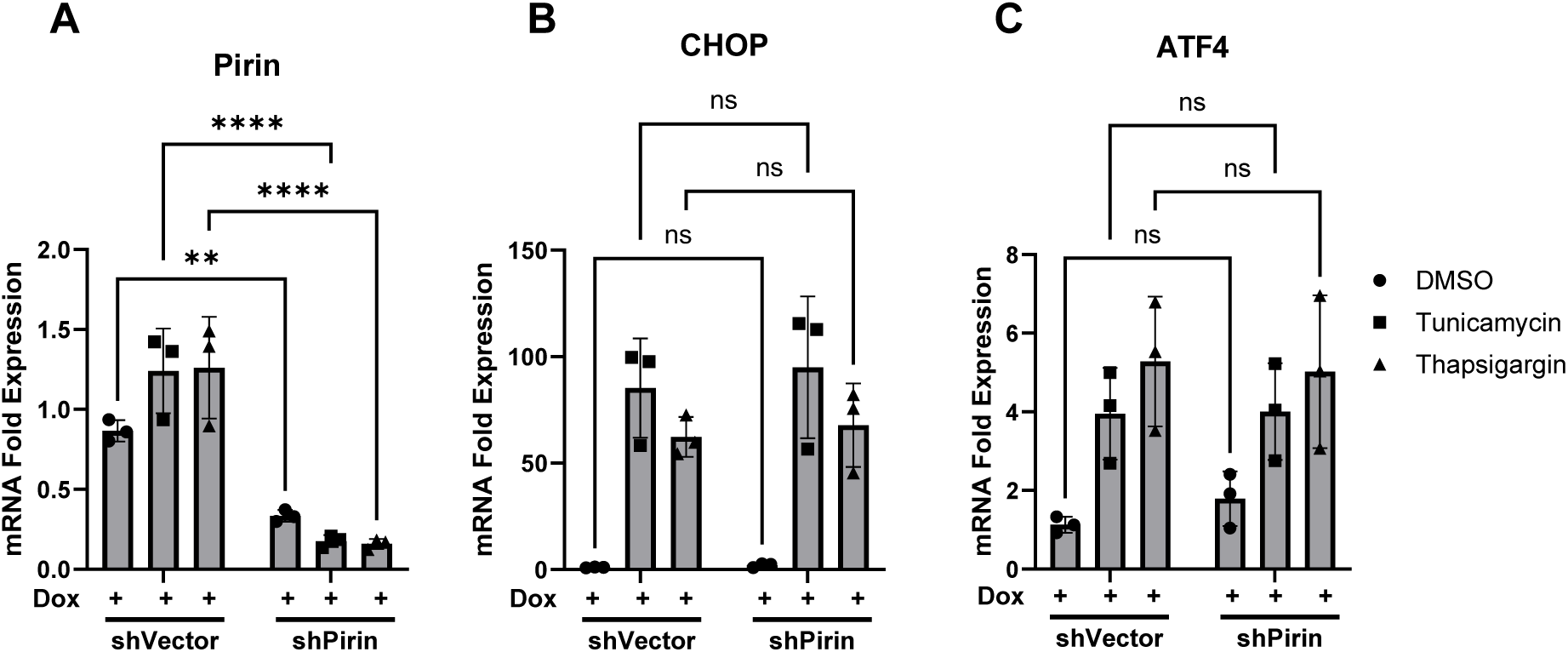
Expression of ER stress genes. CHOP (A) and ATF4 (B) mRNA in pirin knockdown cells (shP+Dox) and control cells (shVector+Dox) was measured by qRT-PCR. ER stress in cells was induced for 5 hours with tunicamycin or thapsigargin before RNA was isolated. Pirin knockdown in shP+Dox cells was confirmed through qRT-PCR (A). Statistical analysis was done by one-way ANOVA in GraphPad Prism 10 with significance values of * = p<0.05, ** = p<0.01, and *** = p<0.001. Error bars show standard deviation.

## Discussion

The aim of this work was to confirm the major proposed mechanism of pirin function (i.e. binding to p65 and regulating NFκB1-mediated gene expression) and then to explore the structural basis of this process. Surprisingly, we were unable to confirm three previous reports (Liu *et al*., 2013; Carrillo-Beltrán *et al*., 2020; Shen *et al*., 2024) that pirin directly binds to or regulates the p65 subunit of NFκB. Despite use of multiple biochemical methods, we found no evidence that pirin binds to the p65 dimer in complex with DNA. Also, by use of pirin knockout and knockdown systems, we found no evidence for pirin modulation of NFκB-mediated gene transcription. We were, however, able to confirm the quercetinase activity of pirin and extend that result to demonstrate a role of pirin in regulation of cellular levels of the flavonol quercetin.

Given the surprising result that pirin didn’t bind, p65 (a.k.a. RelA) in our preliminary SPR studies with nM protein concentrations, we used two additional methods to confirm the lack of binding (gel filtration with 20-40 μM concentrations of purified Fe(III)-pirin and p65 proteins and a fluorescence polarization method to assess pirin’s effect on p65 binding to DNA). Neither showed evidence of a pirin-p65 interaction or modulation of the p65 DNA interaction.

In addition to studies of purified proteins, we also assessed the effect of pirin on cellular canonical NFκB signaling. Using a regulated shPirin knockdown cell line and cells from a novel pirin knockout mouse model, we saw no effect of the lack of pirin on TNFα-induced gene transcription. Furthermore, in our hands, pirin is cytoplasmic with some evidence for ER localization. Given that nearly every paper published about pirin’s biological actions cites early work stating the pirin is nuclear and regulates p65 NFκB function, we felt that it was important to get our “negative” data into the literature.

As our work was being completed, Ma et al, (Ma *et al*., 2024) reported that pirin may be involved in non-canonical NFκB2 signaling in colorectal cancer rather than affecting p65 as stated in most reports on pirin function. They showed co-immunoprecipitation of pirin and RelB when overexpressed in HEK293 cells. Also, pirin was able to disrupt the complex of p52/RelB with a FAS promoter DNA construct. They also hypothesized that the pirin interaction with RelB was in the cytoplasm rather than the nucleus (Ma *et al*., 2024).

Beyond effects of pirin on p65, it has been shown that pirin degrades quercetin and other flavanols (Adams and Jia, 2005; Guo *et al*., 2019). We were able to confirm pirin’s quercetinase activity in biochemical assays and show for the first time that endogenous pirin regulates cellular levels of quercetin. Pirin knockdown leads to higher flavonol levels, especially with low concentrations of quercetin added to the cell cultures. Since flavanols are protective against ROS and oxidative stress (Victor *et al*., 2016; Luo *et al*., 2022), it seems likely that pirin knockdown or pharmacological inhibition could enhance the well-reported beneficial actions of quercetin and other flavonols (Feng *et al*., 2024; Kim *et al*., 2024; Saikia *et al*., 2024).

Due to pirin’s close localization with the ER and pirin’s potential role as a redox-sensor in cells, we tested pirin’s effect on ER stress. ER stress can be caused by altered redox homeostasis in the ER (Cao and Kaufman, 2014) and therefore a connection between pirin and ER stress would have been easily explainable. However, we did not find any changes in ER stress response genes in shPirin cells and could therefore not provide any evidence of pirin’s involvement in ER stress regulation.

Multiple compounds with profound biological effects (TphA, CCT251236, CCG-257081 and related compounds) bind pirin with high affinity, so this work is relevant to their pharmacological mechanisms. The pirin-binding compounds TphA, CCT251236, CCG-257081, and related compounds reduce cancer cell proliferation, induce apoptosis, and synergize with known anticancer agents in melanoma, colorectal cancer, etc. (Miyazaki *et al*., 2010; Cheeseman *et al*., 2016; Evelyn *et al*., 2016; Foda and Neubig, 2023; Ma *et al*., 2024). They also have beneficial effects in *in vivo* cancer models by reducing metastasis, mitigating drug resistance, and enhancing the effects of immunotherapy (Haak *et al*., 2017; Lionarons *et al*., 2019; Foda and Neubig, 2023). They also suppress myofibroblast activation and fibrosis (Sandbo *et al*., 2009; Haak *et al*., 2014a; Bialik *et al*., 2019; Lisabeth *et al*., 2019; Ó hAinmhire *et al*., 2019).

The various mechanisms examined here may play an important role in the compounds’ effects in cancer and fibrosis. Our data strongly suggest that p65 NFκB1-mediated inflammation is not relevant to the actions of the compounds. However, modulation of flavonol levels could play a role as flavonols are actively being explored in both cancer and inflammatory disease therapies (Aghababaei and Hadidi, 2023; Dubey *et al*., 2024; Williamson and Clifford, 2024). Flavonols, such as quercetin and kaemferol are detectable at nM concentrations in plasma of subjects on a normal diet and correlate with predicted dietary flavonol content (Radtke *et al*., 2002). These are two of the best substrates for degradation by pirin (Guo *et al*., 2019) so the pirin-binding compounds could enhance effects of either dietary or therapeutic quercetin and other flavonols. The recent work on the role of pirin to suppress apoptosis and ferroptosis (Hu *et al*., 2021; Ma *et al*., 2024) may represent another key contributor to the actions of pirin inhibitors in cancer.

In summary, our results show that the primary proposed mechanism of biological effects of the protein pirin, modulation of p65/NFκB1 function, could not to be confirmed. Furthermore, pirin’s cellular localization is not primarily in the nucleus in fibroblast-type cells so mechanistic analysis should consider non-nuclear processes. We do confirm that pirin degrades the flavonol quercetin and show for the first time, that pirin-binding compounds (Cheeseman *et al*., 2016; Lisabeth *et al*., 2019), can inhibit this enzymatic activity and increase flavonol levels in cells exposed to exogenous flavonols. These results have implications for better understanding pirin biology and for sorting out potential mechanisms of compounds targeting the protein pirin in cancer and inflammatory diseases.

## Acknowledgements

The authors thank Drs. Elena Demireva and Huirong Xie in the MSU Transgenic and Genome Editing Facility for their assistance in the design and generation of the pirin knockout mouse line.

## Data Availability Statement

The authors declare that all the data supporting the findings of this study are available within the paper and its Supplemental Data.

## Author Contributions

*Participated in research design:* Lisabeth, Meschkewitz, Neubig *Conducted experiments:* Cab Gomez, Leipprandt, Lisabeth, Meschkewitz *Performed data analysis:* Cab Gomez, Lisabeth, Meschkewitz, Neubig *Wrote or contributed to the writing of the manuscript:* Lisabeth, Meschkewitz, Neubig

## Footnotes

*This work was supported by National Institute of Health National Institute of General Medical Sciences [Grant R01GM115459] (RRN) and American Heart Association Predoctoral Fellowship [Grant 23PRE1019204] (MM).

## Abbreviations

ATF4: Activating Transcription Factor 4
BiP: Immunoglobin Binding Protein
BSA: Bovine Serum Albumin
β-Me: beta-Mercaptoethanol
CHOP C/EBP: homologous protein
DMEM: Dulbecco’s Modified Eagle Medium
DTT: Dithiothreitol
EDEM: ER-degradation-enhancing-a-mannosidase-like protein
ER: Endoplasmic Reticulum
FAM: Fluorescein amidite
FP: Fluorescence Polarization
GAPDH: Glyceraldehyde 3-phosphate dehydrogenase
GRP94: Glucose Regulated Protein 94
IL-6: Interleukin 6
IPTG: isopropyl b-D-1-thiogalactopyranoside
IκBα: Inhibitor of kappa B alpha
MEF: Mouse Embryonic Fibroblast
NBCS: New Born Calf Serum
P65RHR: P65 Rel Homology Region
PDI: Protein Disulfide Isomerase
SEC: Size Exclusion Chromatography
SPR: Surface Plasmon Resonance
sXBP1: spliced X-box binding protein 1 total
XBP1: total X-box binding protein 1
TGF-β: Transforming Growth Factor β
TNFα: Tumor Necrosis Factor alpha
usXBP1: unspliced X-box binding protein 1

