## Supplemental Data for "Pirin does not bind to p65 or regulate NFκB-dependent gene expression but does modulate cellular quercetin levels"

**Supplemental Materials**

**Figure S1**

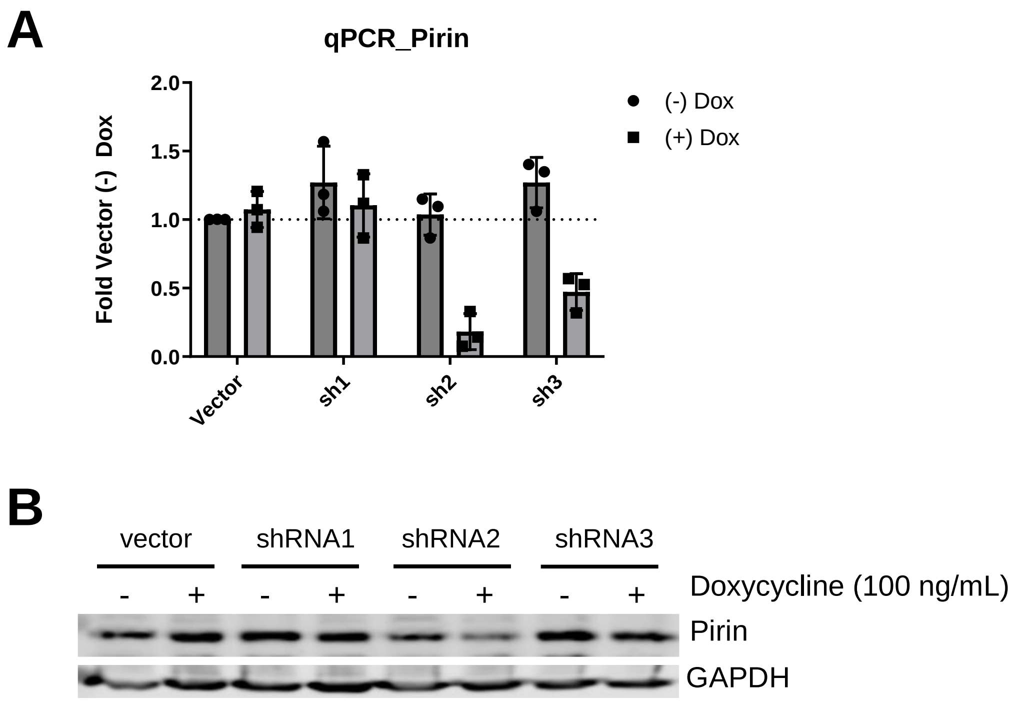


**Figure S1. Generation of knockdown pirin cell lines.** (A) qPCR and (B) Westernblot for pirin expression from 3T3 cells stabely transfected with different silencing RNAs for Pirin.

**Figure S2**

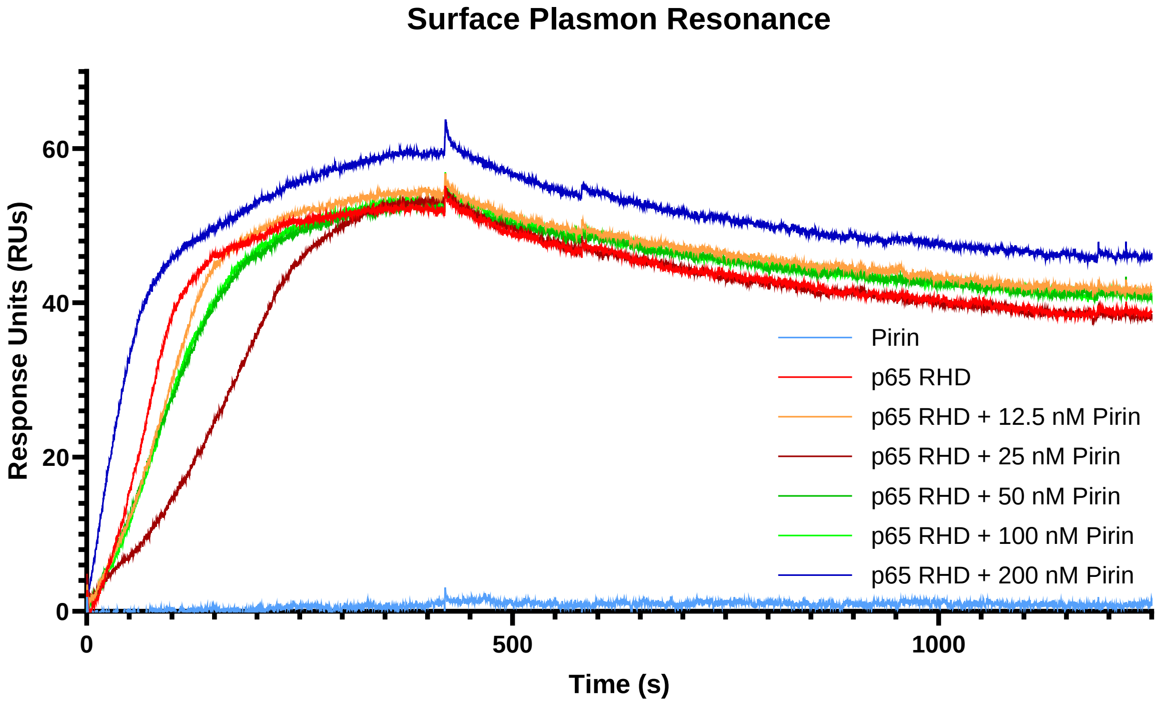


**Figure S2. Surface Plasmon Resonance of p65RHR and pirin.** Biotinylated IκB DNA was bound to Channel A on a Streptavidin Chip. Non-specific DNA was bound to Channel B. P65RHR alone (50 nM) or with different concentrations of Pirin (12.5 – 200 nM), or pirin alone, were run over the chip. Signal from channel B was subtracted from Channel A to correct for non-specific binding. Graph is representative of 3 different experiments.

#### Figure S3

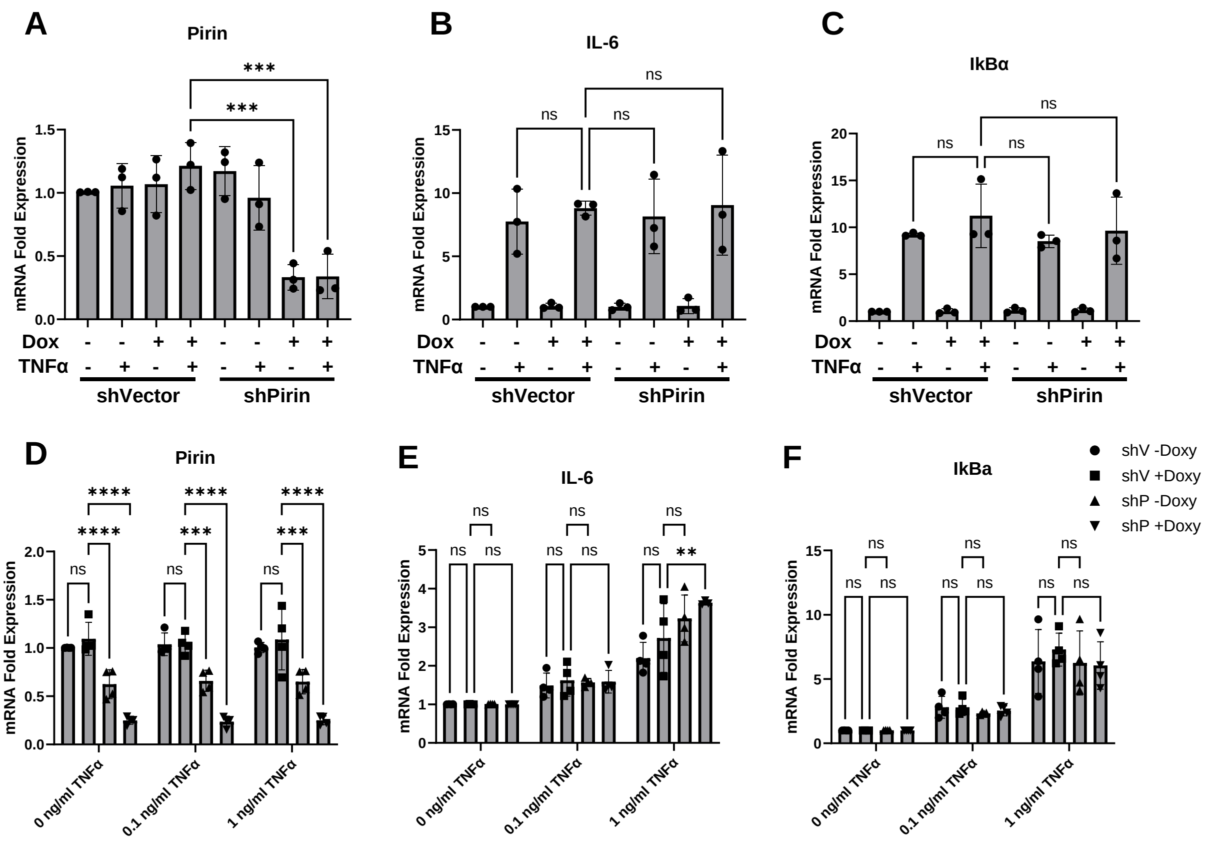


**Figure S3. Expression of NF-kB response genes.** (A) Full dataset shown for Figure 3. Cells were treated with 10 ng/ml TNFα. (B) shVector and shPirin cells treated with different concentrations of TNFα. Data are mean ± SD from 3 independent experiments each with 3 technical replicates. Statistical analysis was done by One-way ANOVA (A-C) or two-way ANOVA with Dunnetts (D-E) in GraphPad Prism 10 with significance values of * = p<0.05, ** = p<0.01, and *** = p<0.001.

**Figure S4**

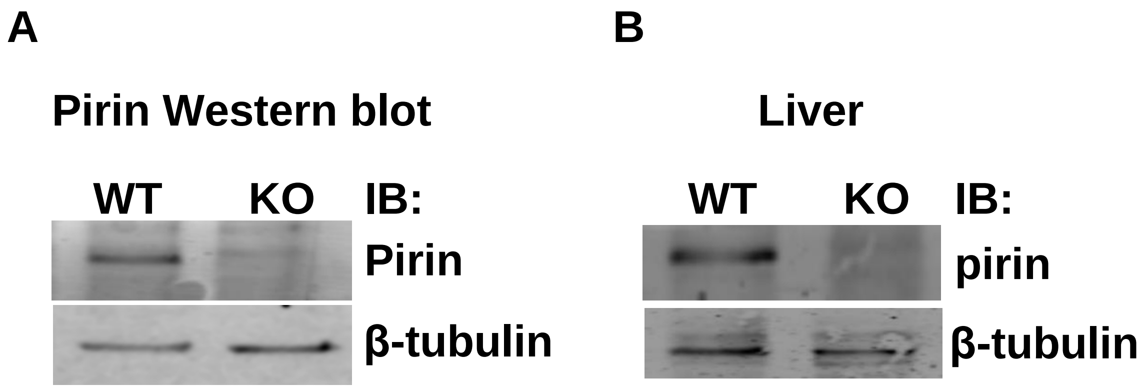


**Figure S4. Pirin knockout mice show global knockout of Pirin.** Westernblots show pirin protein expression in fibroblasts (A) and liver (B) of pirin knockout mice. Beta-tubulin is used as loading control.

**Figure S5**

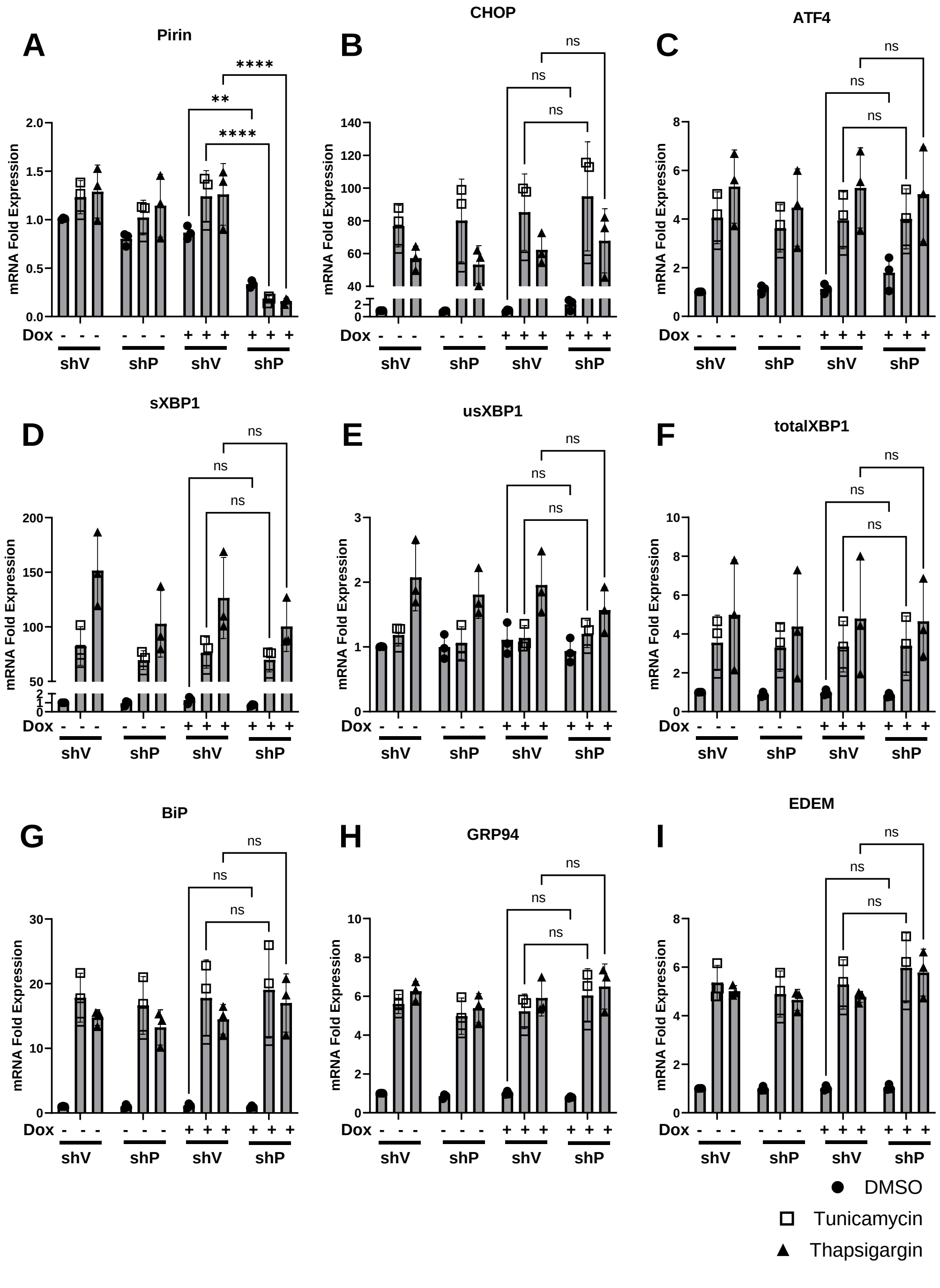


**Figure S5. Expression of ER stress genes** in pirin knockdown cells (shP+Dox) and control cells (shVector-Dox, shVector+Dox, shP-Dox) measured through qRT-PCR. Full data for Figure 5. ER stress in cells was induced for 5 hours with Tunicamycin or Thapsigargin before RNA was isolated. Pirin knockdown in shP+Dox cells was confirmed through qRT-PCR (A). Statistical analysis was done by One-way ANOVA in GraphPad Prism 10 with significance values of * = $p<0.05$, ** = $p<0.01$, and *** = $p<0.001$. Error bars show standard deviation. Genes tested include CHOP (B), ATF4 (C), sXBP1 (D), usXBP1 (E), totalXBP1 (F), BiP (G), GRP94 (H), and EDEM (I).

### Figure S6

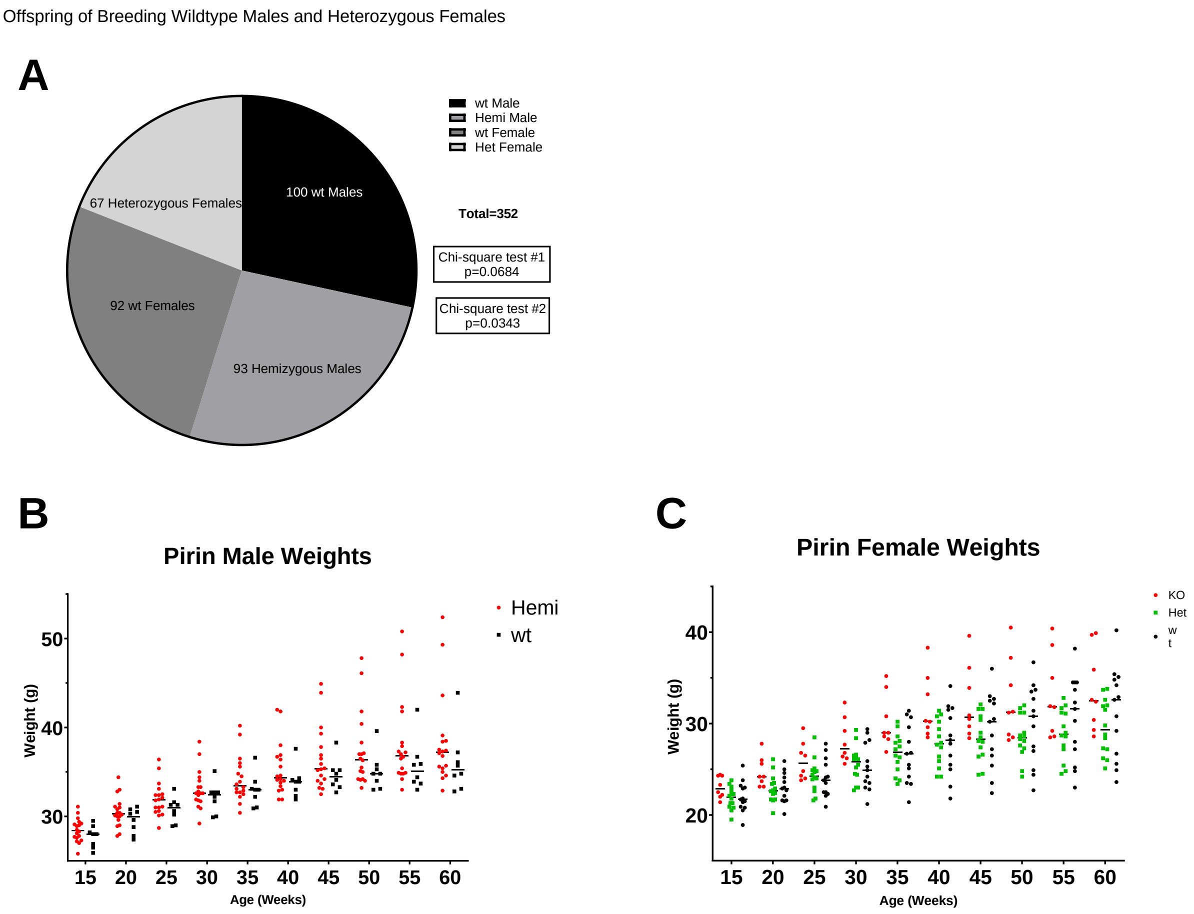


**Figure S6. Breeding outcomes of Pirin knockout mice.**
